# Feature-coding transitions to conjunction-coding with progression through human visual cortex

**DOI:** 10.1101/085381

**Authors:** Rosemary A. Cowell, John T. Serences

**Affiliations:** Department of Psychological and Brain Sciences, University of Massachusetts, Amherst, MA 01003, USA; Department of Psychology, University of California, San Diego, La Jolla, CA 92093-0109, USA; Neurosciences Graduate Program, University of California, San Diego, La Jolla, CA 92093, USA

**Keywords:** visual cortex, feature, conjunction, neuroimaging, object representations

## Abstract

Identifying an object and distinguishing it from similar items depends upon the ability to perceive its component parts as conjoined into a cohesive whole, but the brain mechanisms underlying this ability remain elusive. The ventral visual processing pathway in primates is organized hierarchically: Neuronal responses in its early stages are sensitive to the manipulation of simple visual features whereas neuronal responses in subsequent stages are tuned to increasingly complex stimulus attributes. It is widely assumed that feature-coding dominates in early visual cortex whereas later visual regions employ conjunction-coding in which object representations are different from the sum of their simple-feature parts. However, no study has demonstrated that putative object-level codes in higher visual cortex cannot be accounted for by feature-coding and that putative feature-codes in regions prior to ventral temporal cortex are not equally well characterized as object-level codes. Thus the existence of a transition from feature- to conjunction-coding in visual cortex remains unconfirmed, and, if a transition does occur, its location remains unknown. By employing multivariate analysis of human functional imaging data, we measure both feature-coding and conjunction-coding directly, using the same set of visual stimuli, and pit them against each other to reveal the relative dominance of one versus the other throughout cortex. We provide the first demonstration of a transition from feature-coding in early visual cortex to conjunction-coding in both inferior temporal and posterior parietal cortices. This novel method enables the use of experimentally controlled stimulus features to investigate population-level feature- and conjunction-codes throughout human cortex.

## Significance Statement

Models of object recognition assume that feature-coded representations in early visual cortex are transformed by higher visual regions into an object-level code in which the whole is different from the sum of its parts, but direct evidence for this transformation is lacking. Cowell and Serences use a novel analysis of neuroimaging data to assess stimulus representations throughout human visual cortex, revealing a transition from feature-coding to conjunction-coding along both ventral and dorsal pathways. Representations in occipital cortex contain more information about spatial frequency and contour than about conjunctions of those features, whereas predominantly conjunction-specific information is observed in occipito-temporal, inferotemporal and superior parietal sites. The novel method will enable investigation of feature- and conjunction-coding in other sensory and motor modalities.

## Introduction

Object perception is underpinned by a hierarchical series of processing stages in the ventral visual pathway (1–4). At each successive stage from primary visual cortex (V1) to anterior inferotemporal (aIT) cortex, the visual complexity of the optimal stimuli increases: neurons in V1 are tuned to simple stimulus attributes such as orientation (1, 5); neurons in V4 and posterior inferotemporal cortex (pIT) are selective for moderately complex stimulus features (2, 6); and neurons in aIT prefer partial or complete views of complex objects (2, 7, 8). Data from functional magnetic resonance imaging (fMRI) in humans corroborate these findings: the blood oxygenation level-dependent (BOLD) signal exhibits selectivity for orientation, spatial frequency and color in early visual regions (9–12), but is sensitive to object-level visual properties such as global contour or object category identity in higher regions (13–18). It is widely assumed that the downstream, object-specific representations are constructed through combination of the simple feature representations upstream, but the manner in which this combination occurs remains unknown.

There are at least three possible combination schemes. The first assumes that downstream object-level representations perform 'and-like' operations on upstream feature representations (19), transforming the feature-code into conjunction-sensitive representations in inferotemporal (IT) cortex. This *feature-to-conjunction transition* scheme is assumed by many formal models of object processing (20–27) and accords with electrophysiological data suggesting that IT neurons are selective for complex objects (2, 7, 28). However, when tested with large sets of stimuli many IT neurons show broad tuning, responding to multiple complex objects (7, 29, 30). It is thus possible that apparent object-level selectivity in an IT neuron tested with a small stimulus set could in fact be selectivity for a low-level feature possessed by only a few objects in the set. Thus the data do not rule out a second possible scheme: a *global feature-coding hypothesis*, in which simple features are coded as separate entities in visual cortex and are bound by the synchronization of neural activity rather than by convergence onto a cortically localized representation of the conjunction (31–33). Finally, a third possible coding scheme is a *global conjunction-coding hypothesis*, in which all stations in the hierarchy bind information about simple features together non-linearly to produce conjunctions (34, 35). Under this scheme, the apparent feature-selectivity of neurons in early visual cortex belies a neural code that is optimized for discriminating more complex objects that contain those features. Supporting this account, several studies have reported coding of simple conjunctions of features such as color, form, motion and orientation, in early visual regions (36–41) and coding of more complex conjunctions in both early *and* higher-level visual regions (42).

In order to differentiate between the three alternative representational schemes, we must measure not just the presence of feature-coding or conjunction-coding, but the relative contribution of each, throughout visual cortex. Several recent fMRI studies have examined conjunction-coding in human cortex (38, 39, 42–44). Two of these revealed conjunction-coding for combinations of static visual features in the ventral pathway (39, 42), but neither was able to definitively rule out an explanation of the observed conjunction-code in terms of feature-coding combined with saturation of the BOLD signal. Moreover, no study to our knowledge has explicitly investigated feature-coding - that is, the extent to which neural representations are *more informative* about individual features than their conjunctions - in order to rule out the possibility that apparent feature-selectivity belies an object-level code containing feature information. Without a simultaneous assessment of directly comparable measures of feature-coding and conjunction-coding, the evidence for a transition from one to the other cannot be assessed. Therefore three important questions remain unresolved: Is there evidence for conjunction-coding in the human ventral visual stream when an explanation in terms of BOLD signal saturation is ruled out? If yes, is conjunction-coding dominant even early in the ventral stream or does it emerge with progression along the pathway? If it emerges, at what point does the transition from feature- to conjunction-coding occur?

Using fMRI in humans, we devised a novel stimulus set and multi-variate pattern analysis (MVPA) technique to pit feature-coding against conjunction-coding, thereby measuring the relative prevalence of each throughout visual cortex. Motivated by evidence that the integration of contour elements into global shape (45) and local image features into global texture (46, 47) are key mechanisms by which the ventral pathway constructs complex object representations, we created novel object stimuli by building conjunctions from binary features defined by contour and spatial frequency. These stimuli allowed each cortical region to be probed for information at two levels: features or conjunctions. Feature-information and conjunction-information were assessed throughout cortex using the same neuroimaging dataset and placed in a ratio, allowing direct comparison of the two coding schemes in each cortical region.

## Results

Participants (n=8, two scan sessions each) viewed visual stimuli constructed hierarchically from four binary features to give sixteen unique, conjunctive objects. We used a support vector machine (SVM) to classify patterns of BOLD responses evoked by the stimuli. For each session in each subject, we constructed four two-way feature-level SVM classifiers (one classifier for each binary feature) and one sixteen-way object-level SVM classifier (Figure 1). This yielded both feature- and object-level classification accuracy for a given region of interest (ROI). We next constructed a *Feature Conjunction Index* (FCI) for each ROI by comparing the output of the feature- and object-level classifiers (Figure 1; see *Experimental Procedures*). A positive FCI indicates that the ROI contains more information about individual objects than is predicted from the information it contains about the separate object features, suggesting that the activation pattern is modulated by the presence or absence of specific objects rather than by individual features. A negative FCI indicates that the ROI contains more information about individual features than about whole objects, suggesting that voxel activations are primarily modulated by individual feature dimensions rather than by whole-object identity. FCI values near zero, provided classifier performance is above chance, suggest a hybrid code in which some voxels exhibit feature-coding and others object-coding. Simulation and analysis of synthetic data constructed according to a feature-code, a conjunction-code and a hybrid-code yielded FCI values confirming these intuitions (see *Supporting Information*). Thus, the FCI of a given cortical region indicates the extent to which the voxel-level population code is relatively dominated by a feature-code (negative FCI) or a conjunction-code (positive FCI).

**Figure 1.**
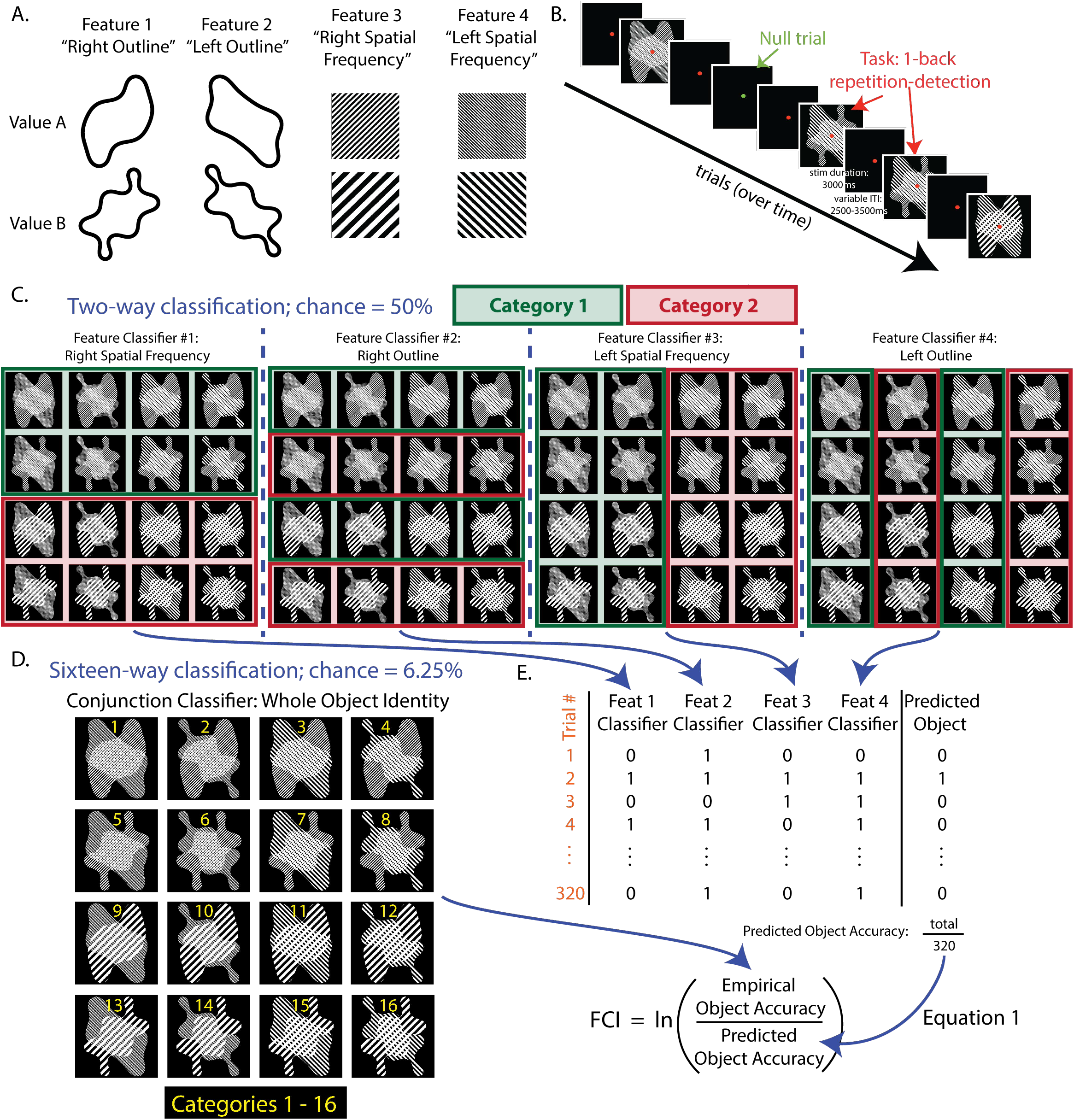
Stimulus Construction, Task Protocol and Multivariate Pattern Classifiers. A. Stimulus Construction. The four binary features from which the 16 object-level stimuli are composed (full stimulus set shown in panel D). Each feature has two possible values: A and B. B. Task Protocol. In each scan session, participants completed 10 experimental runs, each lasting 264 seconds (44 trials of 6 sec duration). A run contained 34-36 stimulus trials (two presentations each of the 16 stimuli in the set, ordered pseudo-randomly, in addition to 2-4 stimuli chosen pseudo-randomly from the set and inserted to create immediate repeats) and 8-10 nulls trials. On stimulus presentation trials, the fixation point was red, stimulus duration was 3sec and the inter-stimulus interval varied between 2.5 and 3.5sec; participants performed a 1-back repetition detection task. On null trials, the fixation point changed to green; participants indicated by button press when they detected a slight dimming of the fixation point, which occurred once or twice per null trial. Participants also completed several sessions of visual search training between the two scans, but we detected no effect of training on our measure of feature- and conjunction-coding (FCI) in cortex (see Supporting Information). See Table S5. C. Feature Classification. Four separate feature classifiers were trained, one for each binary feature defined in the stimulus set. The four feature classification problems are shown in the four panels (Right Spatial Frequency, Right Outline, Left Spatial Frequency, Left Outline) in which the designation of stimuli to feature categories is indicated with red and green boxes. Classifiers used a support vector machine trained with hold-one-out cross-validation (see Supporting Experimental Procedures). See also Tables S1-S2. D. Object Classification. A single object-level classifier was trained to classify the stimuli into 16 categories, each corresponding to a unique stimulus. See Table S3. E. Calculation of the Feature Conjunction Index (FCI). The product of the four feature-level accuracies was used to predict - independently for each trial - the accuracy of a hypothetical object-level classifier whose performance depends only on feature-level information. On each trial, the four feature-classifier responses (defined as 0 or 1 for incorrect or correct) were multiplied to produce a value of 0 or 1 (incorrect or correct) for the hypothetical object-level classifier. Next, the empirically observed object-level classifier accuracy (derived from the sixteen-way conjunction classifier) and the hypothetical object-level accuracy (predicted from the four feature classifiers) were averaged over trials and placed in a log ratio (Equation 1). Whenever the empirically observed object classifier accuracy exceeds the hypothetical object accuracy predicted from feature classifier accuracies, FCI is positive. Whenever the feature classifier accuracies predict better object-level knowledge accuracy than is obtained by the object classifier, FCI is negative. See Figure S1.

### A Transition from Feature- to Conjunction-Coding in Occipito-Temporal Cortex

We investigated whether early visual cortex employs feature-coding or conjunction-coding, and whether one scheme transitions to the other with progression toward the temporal lobe, by examining a series of functionally or anatomically defined ROIs: V1, V2, V3 and lateral occipital cortex (LOC) (Figure 2). Taking stimulus-evoked activation patterns from across each ROI, we trained the classifiers using hold-one-out cross-validation (see *Supporting Experimental Procedures*) and computed FCI for each subject and session separately. FCI differed significantly across ROIs (F(5,35) = 15.78, p < 0.001, η^2^=0.693), but not across sessions (F(1,7) = 0.004, p = 0.95, η^2^=0.001), with no interaction between Session and ROI (F(5,35)= 0.133, p = 0.984, η^2^=0.019). Ordering the ROIs as [V1, V2v, V2d, V3v, V3d, LOC] revealed a significant linear trend (F(1,7) = 38.14, p < 0.001; η^2^= 0.845), indicating a transition from feature-coding toward conjunction-coding along early stages of the visual pathway. Regions V1 through V3 exhibited negative FCIs indicative of feature-code dominance, while LOC revealed a numerically positive FCI that was significantly greater than in all other ROIs (revealed by non-overlapping 95% CIs, Figure 2), suggesting a hybrid code with substantial degree of conjunction-coding.

**Figure 2.**
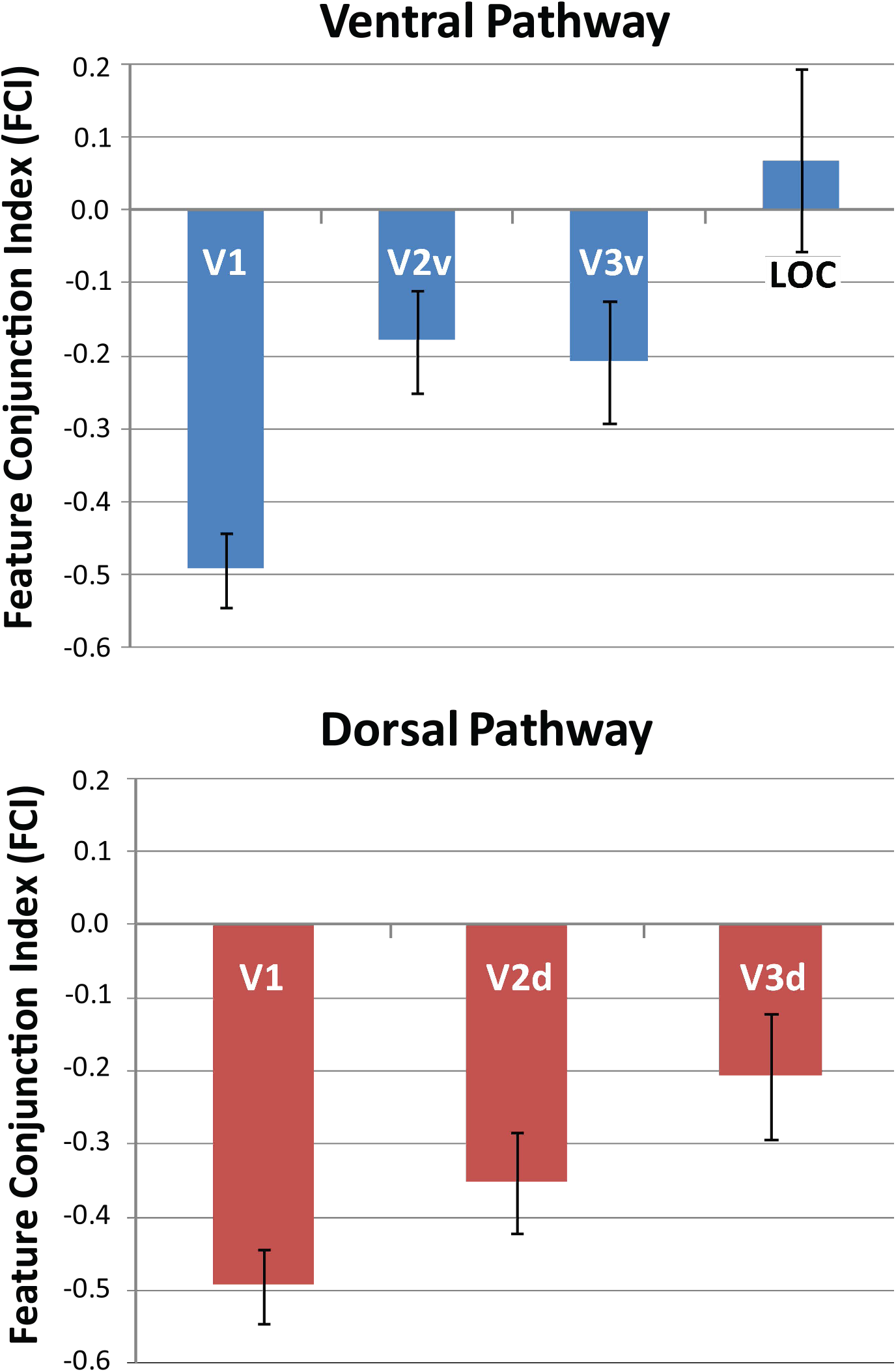
Feature Conjunction Indices (FCI) derived from ROI-based analyses. Mean FCI (averaged over two sessions in each subject) for ROIs in early ventral visual stream (left panel: V1, V2v, V3v, LOC) and early dorsal stream (right panel: V1, V2d V3d). V1 is duplicated in left and right plots for ease of comparison. FCI is the natural logarithm of the ratio of object classifier accuracy to the product of the four feature classifier accuracies (see Figure 1 and Supporting Experimental Procedures). Positive FCI reflects conjunction-coding; negative FCI reflects feature-coding. Error bars show 95% Confidence Intervals (CIs) determined by within-subjects bootstrap sampling with replacement over 10,000 iterations. n = 8 subjects, 2 scan sessions each. See Table S4.

### Feature- and Conjunction-Coding Throughout Visual Cortex

In order to assess feature- and conjunction-coding in cortical representations of objects beyond region LOC, and in the dorsal visual pathway, we examined all of visual cortex using a searchlight approach (48). At each spherical ROI, we performed classifier analyses (using hold-one-out cross-validation and screening out spheres in which classifier accuracy did not exceed chance, see *Supporting Experimental Procedures*) and computed FCI, mapping the FCI value back to the centroid voxel of the sphere. In the group-averaged FCI map (Figure 3), occipital regions exhibit the most negative FCIs (green) indicative of feature-coding. With progression into regions anterior and superior to the occipital pole, FCI values first become less negative (blue) – suggesting transition to a hybrid code – and then become positive (orange/yellow) in occipito-temporal and posterior parietal regions, indicating the emergence of conjunction-coding in both ventral and dorsal pathways. Examination of FCI maps in individual subjects revealed the same pattern in every subject in both sessions: strongly negative FCI values in occipital regions with a transition to positive FCI values towards temporal and parietal regions.

**Figure 3.**
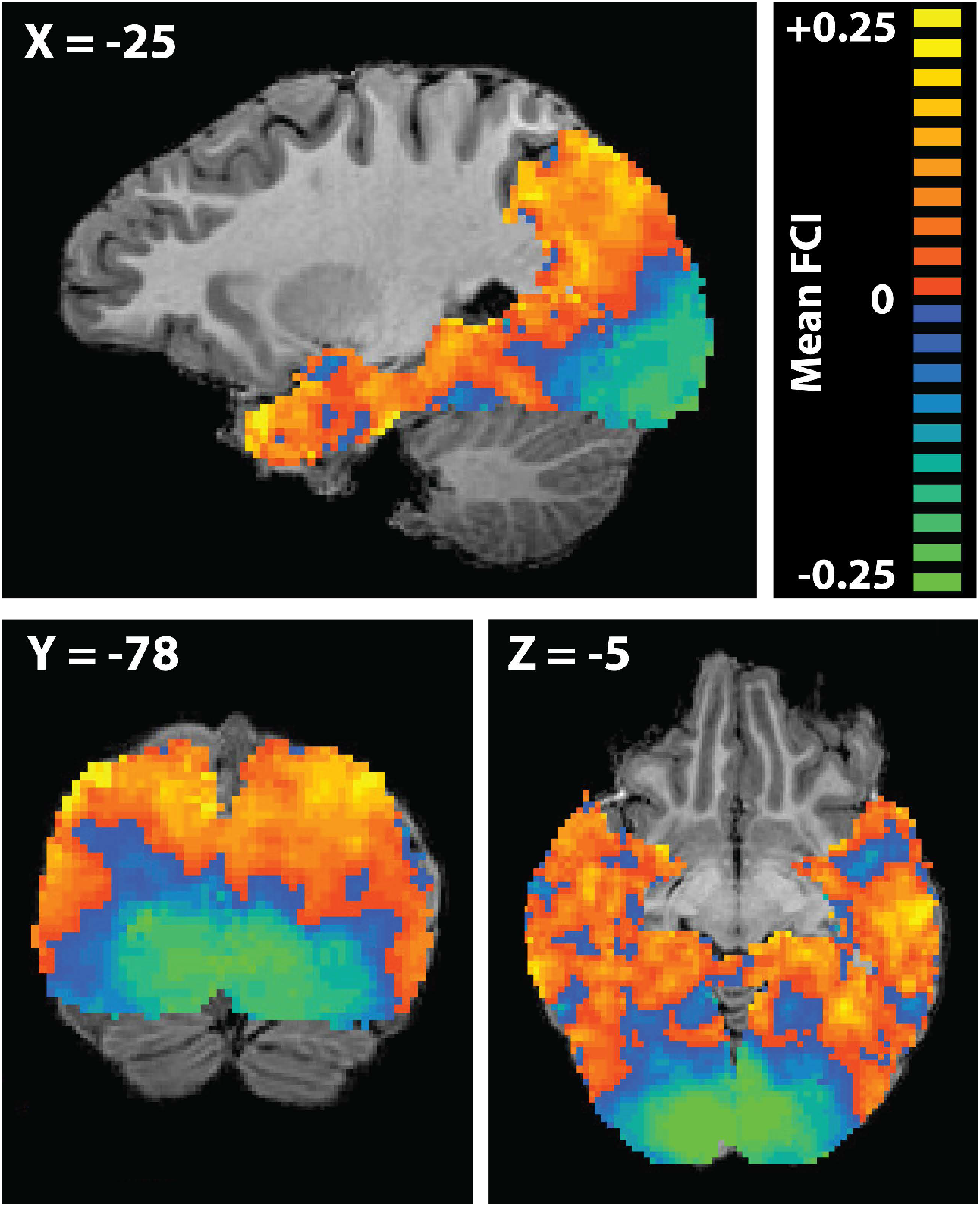
FCI derived from whole-brain searchlight analyses. Group mean FCI produced by a spherelight MVPA analysis assessing conjunction- versus feature-coding throughout visual cortex. A sphere of radius 5 functional voxels was swept through the imaged volume, constrained by a subject-specific grey-matter mask encompassing occipital, temporal and posterior parietal cortex. Taking each voxel in turn as the centroid of a spherical ROI, the feature and object classifiers were trained and their accuracies combined to produce a FCI which was entered into the map at the location of the centroid voxel. Orange indicates positive FCI (conjunction-coding), blue indicates negative FCI (feature-coding). Centroid voxels for which classifier performance did not exceed chance, as determined by a binomial test, were removed from individual subject maps (see Supporting Experimental Procedures). FCI maps were constructed for each subject and scan session individually, then spatially smoothed using a Gaussian kernel (FWHM=2 functional voxels). Smoothed maps were averaged across two sessions for each subject, and across subjects. Scale is truncated at ±0.25 for optimal visualization of the data; some voxels possessed FCI values > +0.25 or < -0.25.

### Quantifying the Transition from Feature- to Conjunction-Coding

Next, we sought to quantify the relationship between cortical location and FCI in both ventral and dorsal pathways. To do so, we devised a metric to specify the location of the voxels in each pathway by defining three vectors in Talaraich co-ordinates: a '*Posterior Ventral*' vector with its origin in the occipital pole extending to the center of LOC; an '*Anterior Ventral*' vector with its origin in LOC extending to the anterior tip of the temporal pole; and a '*Dorsal*' vector with its origin in inferior posterior occipital cortex extending to the most superior/anterior point of the dorsal pathway in the scanned volume (in Brodmann Area 7). For each vector we defined a bounding box around the vector to constrain the anatomical region from which voxels were drawn (Figure 4; see *Supporting Experimental Procedures*) and projected the Talairach co-ordinates of voxels within the box onto the vector, yielding a scalar value for each voxel that specified its location along the vector. Finally, for each subject, for all three vectors in each hemisphere separately, we computed (i) the correlation between the location of a voxel and the FCI of the spherical ROI centered on that voxel, and (ii) the slope of the best fitting regression line relating voxel location to FCI (Figure 4). The correlation between location and FCI was positive and highly significant in all 8 subjects in both hemispheres for the *Dorsal* vector (p< 0.0001) and the Posterior Ventral vector (p<0.01), reflecting a robust transition from feature-coding at the occipital pole to representations more dominated by conjunction-coding in lateral occipital and superior parietal cortices, respectively. For the *Anterior Ventral* vector, the correlation was positive and significant (p<0.01) in both hemispheres for 5 out of 8 subjects; in 1 subject (yellow in Figure 4) the left hemisphere was negatively correlated (a decrease in FCI with anterior progression; p<0.05) and the right was positively correlated (p<0.001); in the 2 remaining subjects (black and magenta in Figure 4), the left hemisphere was significantly negatively correlated (p<0.0001) and the right was not correlated (p>0.2). The slopes of the best fitting regression lines differed for the three vectors (F(2,14)=23.71, p<.001), but did not differ by hemisphere (F(1,7)=.026, p=.877), with no hemisphere by vector interaction (F(2,14)=2.889, p=.089). Slopes for the *Anterior Ventral* vector were smaller than for the *Posterior Ventral* (p < 0.001) and *Dorsal* (p < 0.0001) vectors, which did not differ from each other (p = 0.62). These results suggest that, for the present stimulus set, the greatest transition from feature- to conjunction-coding occurs in posterior regions, in both ventral and dorsal pathways. A possible reason for the shallower transition toward conjunction-coding in the anterior ventral pathway is that the object-level conjunctions comprising these simple, novel objects may be fully specified in relatively posterior sites, just beyond the occipito-temporal junction (Figures 3 and 5). Indeed, the stimuli with which Erez et al. (42) revealed conjunction coding in anterior temporal regions were 3-dimensional, colored objects that likely better engaged anterior visual regions.

**Figure 4.**
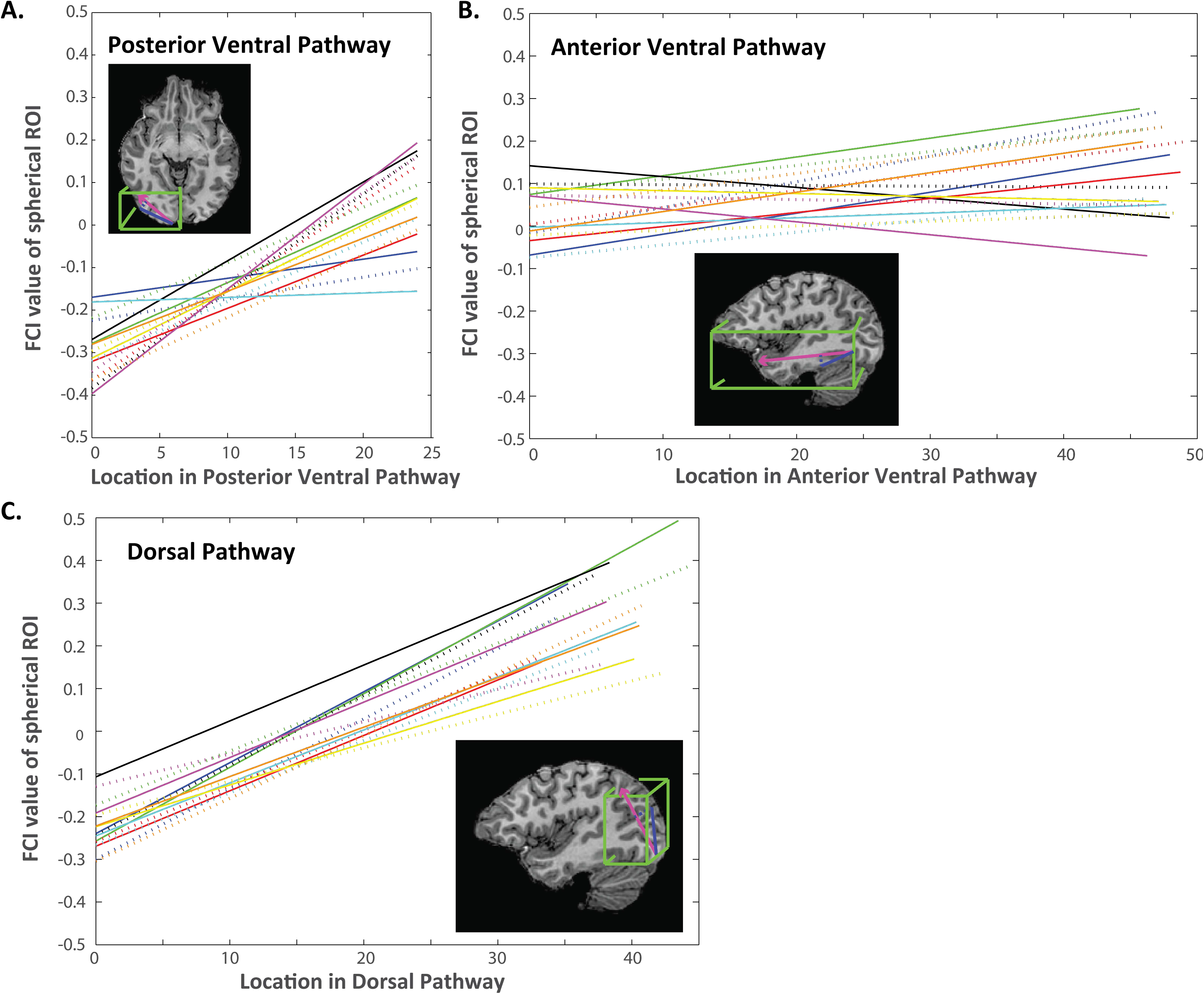
Quantification of the transition from feature- to conjunction-coding in the ventral and dorsal streams. Insets show, in pink, the approximate extent and position of the three defined vectors Posterior Ventral, Anterior Ventral and Dorsal; in blue, the projection of a voxel location onto the vector to derive a scalar value for the voxel position; in green, the bounding box defining the brain region included as part of each pathway (a subject-specific anatomical mask including only grey matter in occipital, temporal and parietal lobes was also applied). Plots show the best fitting regression lines relating the location of a voxel in each of the three pathways to the FCI for the spherical ROI surrounding the voxel. Each line shows one subject in one hemisphere; colors indicate different subjects; solid and dashed lines show Left and Right hemispheres, respectively. The far endpoint of the vector is more distant from occipital cortex (i.e., the vector is longer) for the Dorsal than the Posterior Ventral pathway: this may account for the higher FCI value at the vector endpoint in the Dorsal than the Posterior Ventral pathway, given that regression line slopes in the Dorsal and Posterior Ventral pathways were similar (x and y-axes use the same scale for all 3 plots).

### Cortical Sites of Extreme Feature- and Conjunction-Coding

Finally, to search for cortical sites demonstrating statistically reliable extremes of feature- or conjunction-coding, we compared the group mean FCI at each voxel to zero (two-tailed t-test, False Discovery Rate (FDR) corrected; positive t-values indicate conjunction-coding and negative t-values feature-coding). This analysis assumes anatomical and functional correspondence of points in Talairach space across subjects; it is therefore conservative, particularly for conjunction-coding, the cortical sites of which are likely more widely and variably distributed across subjects. Nonetheless, we revealed a large occipital region of feature-coding along with multiple conjunction-coding sites throughout occipito-temporal, ventral temporal and parietal cortices, extending into the parahippocampal gyrus, medial temporal lobe and anterior temporal pole (Figure 5). In line with a *feature-to-conjunction transition hypothesis*, all conjunction-coding sites were located anterior or superior to the feature-coding sites, which were confined to the occipital lobe (excepting a single more anterior feature-coding voxel at [10–51 5], in the inferior posterior cingulate).

**Figure 5.**
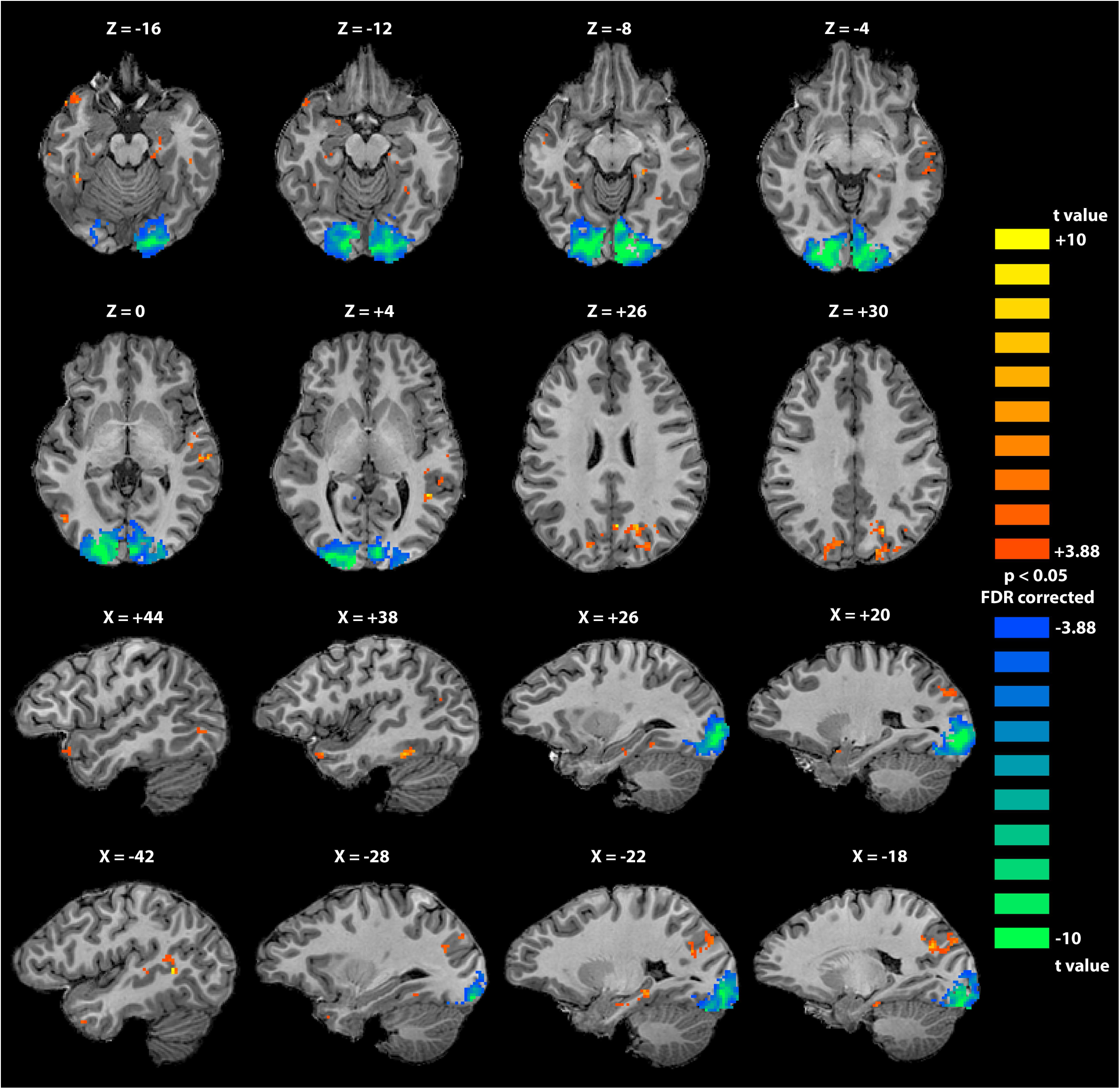
Cortical sites of feature- and conjunction-coding observed at the group level. Statistical map shows the results of a t-test at each voxel comparing the group mean FCI value associated with the spherical ROI surrounding that voxel to zero. The map was thresholded at p=0.05, two-tailed (FDR-corrected for multiple comparisons). Blue voxels possess FCI values significantly less than zero (feature-coding); orange voxels possess FCI values significantly greater than zero (conjunction-coding). All but one voxels with statistically reliable negative FCI values were located in occipital cortex, whereas all voxels with statistically reliable positive FCI values were located anterior or superior to the occipital feature-coding regions. Voxels exhibiting significant conjunction coding appeared in multiple sites bilaterally, including posterior parietal lobe, fusiform gyrus, parahippocampal gyrus (including left perirhinal cortex), hippocampus and the anterior temporal pole. Axial slices are radiologically flipped (left hemisphere appears on the right). In sagittal slices, positive X co-ordinates indicate right hemisphere.

## Discussion

We measured the relative dominance of feature- versus conjunction-coding throughout visual cortex by directly comparing evidence for the two coding schemes in each cortical region. Critically, evidence for both schemes was derived from a common neuroimaging dataset, acquired while participants viewed systematically constructed conjunctive visual stimuli. We revealed a transition from feature-dominated to conjunction-dominated coding, with progression from primary visual cortex into temporal cortex (Figures 2, 3 and 4). This constitutes the first direct evidence for an object representation scheme within the ventral pathway that is characterized by a *transition from feature- to conjunction-coding*. Strikingly, the same shift from feature- to conjunction-coding was evident in the dorsal pathway, where it was as steep and robust as in the posterior ventral pathway (Figure 4). We revealed significant conjunction-coding in a range of sites bilaterally, including posterior parietal cortex, fusiform and parahippocampal gyri, medial temporal lobes and anterior temporal poles (Figure 5). These findings concur with prior reports of conjunction-coding throughout ventral visual stream and perirhinal cortex (42) and in parietal lobe (43, 44). In addition, this study provided the first explicit investigation of feature-coding in human cortex, revealing significant selectivity for features over conjunctions across posterior occipital regions (Figure 5). For the present stimulus set, the transition from feature to conjunction-coding – the zero point of the FCI measure – occurred in the ventral pathway near the occipito-temporal junction, and in the dorsal pathway at approximately the superior border between Brodmann areas 18 and 19 of the occipital lobe (Figure 3).

The finding of a clear transition from feature- to conjunction-coding in the dorsal pathway aligns with the well-documented role of parietal cortex in feature binding (49–53) but also with recent claims that the role of dorsal stream in vision extends beyond spatial processing or attentional binding. That is, our observation of object-specific coding in parietal cortex – elicited while participants discriminated between highly similar visual objects – suggests that the dorsal pathway constructs content-rich, hierarchical representations containing abstract information that is critical for object identification, in parallel with the ventral stream (54, 55).

While the present finding of feature-coding in V2 appears to contradict previous reports of conjunction-coding in this region for stimuli comprising combinations of simple features (36, 39), we suggest that the two sets of results are compatible: negative FCI values in V2 do not rule out the existence of conjunction-based information altogether, they merely imply that any conjunction-code is relatively swamped by a stronger feature-code. A strength of the present method is that it reveals the relative dominance of feature-based versus conjunction-based information.

An analysis of synthetic data supported our interpretations of the FCI metric (see Supporting Information). Across a range of SNR values, provided that classifier performance was above chance, near-zero FCI values were obtained only with data synthesized using a hybrid code and, critically, negative FCIs were produced only by synthetic feature-coded data while positive FCIs emerged only from synthetic conjunction-coded data. Thus, FCI provides a reliable measure of the relative contributions of feature- versus conjunction-coding, and varying noise levels do not produce distortions in FCI that lead to qualitatively erroneous conclusions. Accordingly, the method mitigates greatly against the problem of varying noise levels that has complicated prior attempts to use fMRI to compare the neural code across diverse brain regions. For example, if a standard MVPA method detects greater classification accuracy for simple visual features in early visual cortex than in later visual regions, this could be because the neural representations in early visual regions exhibit stronger feature-coding, or because early visual cortex, which is situated peripherally, emits a BOLD signal with a greater SNR. In the present method, because the feature- and conjunction-coding measures are placed in a ratio, and both measures are affected by the noise in a given cortical region, the relative dominance of feature- versus conjunction-coding in a region maps consistently to negative versus positive FCI values in the face of varying noise levels. We can therefore determine which regions exhibit feature- versus conjunction-coding and approximately localize the point of transition from one to the other in cortex, for a given stimulus set.

Because the binary features used in our dataset included two shape outlines and two spatial frequencies of fill pattern, the intersection of those features created novel shapes and textures; any voxels sensitive to such incidental features may have improved conjunction classification accuracy by reducing the 16-way problem to a 4-way problem (see, for example, the global outline shared by stimuli 1, 3, 9 and 11 in Figure 1D). However, we suggest that while incidental feature-coding may render negative FCIs less negative, it is unlikely to produce positive FCIs in a brain region entirely insensitive to experimenter-defined conjunctions, and unlikely even to produce zero FCIs unless the region overwhelmingly codes for incidental features. As the analysis of synthetic hybrid-coded data demonstrated, activation patterns containing a 50:50 mixture of feature-sensitive and conjunction- sensitive voxels produce FCI values near zero. Any voxels sensitive to incidental features would confer less benefit to the conjunction classifier than truly conjunction-sensitive voxels (since each incidental feature is shared by 4 unique objects), such that a feature-coding ROI would need to contain many more voxels sensitive to incidental features than voxels sensitive to the experimenter-defined binary features for its negative FCI to approach zero. Any ROI containing such a reliable preference for the more infrequent incidental features over the experimenter-defined features would be better characterized as a region that codes for mid-level conjunctions (textures or global outline) than as a feature-coding region. In line with the idea that sensitivity to mid-level conjunctions may contribute to zero FCI values, we observed the zero-point in FCI near to area LOC – known to be sensitive to global shape (13, 15, 18) – and area V4 – known to exhibit texture-selective responses (56–58).

In sum, our novel method permits the systematic investigation of feature- and conjunction-coding, and may be applicable not just to vision but to other modalities such as audition or motor action (59, 60). Within vision, the method enables future investigation of a range of features not included in our stimulus set (including color, orientation and motion), in order to examine how conjunction-coding emerges for different feature types and combinations. The present finding of a transition from feature- to conjunction- coding along both ventral and dorsal visual pathways has implications for theories of the functional architecture of visual object processing.

## Author Contributions

Conceptualization, R.A.C.; Methodology, R.A.C.; Software, R.A.C. and J.T.S.; Formal Analysis, R.A.C. and J.T.S.; Resources, R.A.C. and J.T.S.; Writing – Original Draft, R.A.C.; Writing – Review and Editing, R.A.C. and J.T.S.; Visualization, R.A.C; Funding Acquisition, J.T.S and R.A.C.

## Acknowledgements

We thank David Huber for helpful discussions. This work was funded by NSF CAREER Award 1554871to R.A.C., by R01EY025872 to J.T.S. and by NSF grant BCS-0843773.

